# Morphometric Analysis of Actin Networks

**DOI:** 10.1101/2024.06.10.598236

**Authors:** Oghosa H. Akenuwa, Jinmo Gu, Andreas Nebenführ, Steven M. Abel

**Author notes:** &.

## Abstract

The organization of cytoskeletal elements is pivotal for coordinating intracellular transport in eukaryotic cells. Several quantitative measures based on image analysis have been proposed to characterize morphometric features of fluorescently labeled actin networks. While helpful in detecting differences in actin organization between treatments or genotypes, the accuracy of these measures could not be rigorously assessed due to a lack of ground-truth data to which they could be compared. To overcome this limitation, we utilized coarse-grained computer simulations of actin filaments and crosslinkers to generate synthetic actin networks with varying levels of bundling. We converted the simulated networks into pseudo-fluorescence images similar to images obtained using confocal microscopy. Using both published and novel analysis procedures, we extracted a series of morphometric parameters and benchmarked them against analogous measures based on the ground-truth actin configurations. Our analysis revealed a set of parameters that reliably reports on actin network density, orientation, ordering, and bundling. Application of these morphometric parameters to root epidermal cells of *Arabidopsis thaliana* revealed subtle changes in network organization between wild-type and mutant cells. This work provides robust measures that can be used to quantify features of actin networks and characterize changes in actin organization for different experimental conditions.

## Introduction

Eukaryotic cells are highly ordered systems that maintain their shape and internal organization with the help of the cytoskeleton, a dynamic network of protein filaments that provides both structural integrity and tracks for intracellular transport. The organization of the cytoskeleton, composed of microtubule and actin filament networks, is crucial for normal cell functions, normal growth, and the development of multicellular organisms. In plant cells, the actin cytoskeleton defines the direction of cell expansion (J. Li et al., 2015; Szymanski & Staiger, 2018). In addition, plant actin filaments play a central role during interphase by providing the tracks for organelle movements, which are driven by class XI myosin motors (Nebenführ & Dixit, 2018).

During cytoplasmic streaming, major streams of active organelle movements follow actin filament bundles, and the behavior of individual organelles is strongly influenced by the arrangement of actin filaments (Geitmann & Nebenführ, 2015). In the well-studied pollen system, actin filaments form prominent longitudinal bundles that are required for organelle movements and cell expansion. Any disruption of the actin organization, such as caused by the loss of proteins that affect actin bundling, also disrupts tip growth (Zhang et al., 2019). Interestingly, alterations of the level of actin filament bundling also lead to larger growth defects that go beyond individual cells. For example, loss of two isoforms of the villin actin crosslinker in *Arabidopsis* leads to a lack of growth coordination between neighboring cells, which results in twisted growth of roots, stems, and leaves (van der Honing et al., 2012). Similar effects have been reported for a villin mutant in rice where the twisted growth is accompanied by a faster gravitropic response (Wu et al., 2015). This latter effect is likely caused by faster recycling of the auxin transport protein PIN2, a conclusion that is further supported by the discovery that the auxin transport inhibitor 2,3,5-triiodobenzoic acid (TIBA) triggers villin oligomerization, increased bundling of filaments, and reduced recycling of PIN2 (Zou et al., 2019). Actin filament organization is also involved in pathogen responses. For example, VILLIN3 is a target of the pathogen-associated molecular pattern (PAMP) signaling, resulting in less bundling of filaments and closing of the stomata to prevent entry of pathogens (Zou et al., 2021).

These observations demonstrate that the actin cytoskeleton is an integral component of plant cells. The actin cytoskeleton is also highly sensitive to environmental conditions and can readily adapt to external signals (Yuan et al., 2023). Quantitative image analysis is a key tool that allows one to capture the complex organization of actin filaments and better understand cellular processes involving the cytoskeleton. Over the years, a number of image-based analysis tools have been developed to characterize different actin filament architectures found in both plant and animal cells (Gan et al., 2016; Jacques et al., 2013; Kimori et al., 2016; P. Li et al., 2023; Liu et al., 2022).

A frequently used protocol for image-based analysis of 2-D microscopy images of the plant actin cytoskeleton was developed by Higaki et al. (Higaki et al., 2010). This method, based on confocal images of fluorescently labeled actin filaments, identifies filaments by a combination of filtering, thresholding, and skeletonization (Higaki et al., 2010). The main measures of actin cytoskeletal morphology from this analysis are measures of filament orientation, network density, and network bundling. These morphological measures are capable of capturing biologically relevant and interpretable differences in cytoskeletal organization between different experimental conditions. The largely automated image analysis pipeline is also conducive to rapid analysis of large numbers of images. These approaches were useful, for example, in dissecting the role of the actin cytoskeleton during growth (Henty et al., 2011; Rosero et al., 2013; Scheuring et al., 2016; Takatsuka et al., 2018) and in establishing that actin filaments are rearranged in response to pathogen attack (Henty-Ridilla et al., 2013; Inada et al., 2016; Lu et al., 2020).

While the morphological measures of Higaki et al. have provided valuable insights into the interplay between cellular processes and cytoskeletal organization, there are situations where these published parameters break down. For example, a bundling measure based on the skewness of pixel intensities is sensitive to the overall level of bundling in the cells and the relative signal intensities of the images. In such cases where cells have a high degree of bundling, the coefficient of variation of pixel intensities has been suggested as a more robust measure of bundling (Higaki et al., 2020). Furthermore, these published morphological measures were benchmarked using artificial images that do not resemble cytoskeletal networks in real cells. Given the limitations in imaging the complete actin filament network with optical microscopes, the extent to which the morphological measures proposed by Higaki et al. capture information about the cytoskeletal organization in real cells remains unknown.

Computer simulations based on realistic biophysical interactions provide a means to generate simulated actin networks with single-filament resolution. Such simulated networks offer an opportunity to test the fidelity of morphological parameters against a network where the locations of all filaments are known and ground-truth values of the morphological parameters can be determined. Several simulation frameworks such as AFINES (Freedman et al., 2017), MEDYAN (Popov et al., 2016), and CytoSim (Nedelec & Foethke, 2007) have proven useful in simulating realistic actin networks at biologically relevant length scales. For example, simulations have shown the formation of random mesh-like networks, ordered bundle structures, and contracted network structures similar to those observed both in cells and in reconstituted actin-crosslinker systems. In these simulations, the resulting network structures were regulated by the crosslinker density (Akenuwa & Abel, 2023; Belmonte et al., 2017; Cyron et al., 2013; Freedman et al., 2018; Popov et al., 2016). By applying image analysis methods to realistic simulated networks, measured morphological measures of actin organization can be compared to analogous measures based on the ground-truth configuration of actin filaments. Using this approach, we identify a set of robust morphometric measures of actin filament organization and use them to characterize wild-type and mutant root epidermal cells of *Arabidopsis thaliana*.

## Results

### Simulations and image processing of crosslinked actin networks

We simulated actin filaments in cell-sized regions with varying numbers of crosslinkers (*N_c_*) using the Actin Filament Network Simulation (AFINES) model (Freedman et al., 2017). AFINES is a coarse-grained model that simulates the dynamics of individual semiflexible actin filaments and actin crosslinking proteins. AFINES has been used to study different network architectures, ranging from mesh-like to highly-bundled networks, by modulating the crosslinker density (Akenuwa & Abel, 2023; Freedman et al., 2017, 2018). The simulated actin networks served as ground-truth filament configurations and were used to generate pseudo-fluorescence images analogous to confocal microscopy images. We then processed the pseudo-fluorescence images using a pipeline developed to extract linear features from experimental microscopy images. The processing pipeline consisted of smoothing, background subtraction, and filtering to enhance linear features, followed by thresholding to reveal skeletonized images. The pipeline is analogous to the procedure described in Higaki et al. (2020) and Ueda et al. (Ueda et al., 2010). The skeletonized image was used as a mask, and the pixel intensities along the skeletonized features were used to maintain information about the local number of filaments.

Our simulations revealed a variety of network structures with varying degrees of filament bundling depending on the number of crosslinkers in the system. Figure 1 shows pseudo-fluorescence images and the associated skeletonized images for representative networks with 0, 200, 400, and 800 crosslinkers. The pseudo-fluorescence images highlight the local density of filaments and show the increased degree of bundling as the number of crosslinkers increases. When *N_c_* = 0 and 200, meshwork structures extended across the entire simulation domain. Skeletonized images reproduced a fine meshwork of lines reflecting the meshwork structure in the original images. With *N_c_* = 400, prominent bundles with stray filaments emerged in the network. With *N_c_* = 800, two main bundles spanned the entire domain and tended to align with the long axis of the simulation domain. By aligning with the longer dimension of the confinement area, the filament bundles reduce energetically unfavorable bending that would be required to fit in the shorter dimension. This is consistent with observations from previous studies (Alvarado et al., 2014; Claessens et al., 2006). For bundled networks, skeletonized images mainly capture the pronounced bundle structures and some of the finer filament structures.

**Figure 1:**
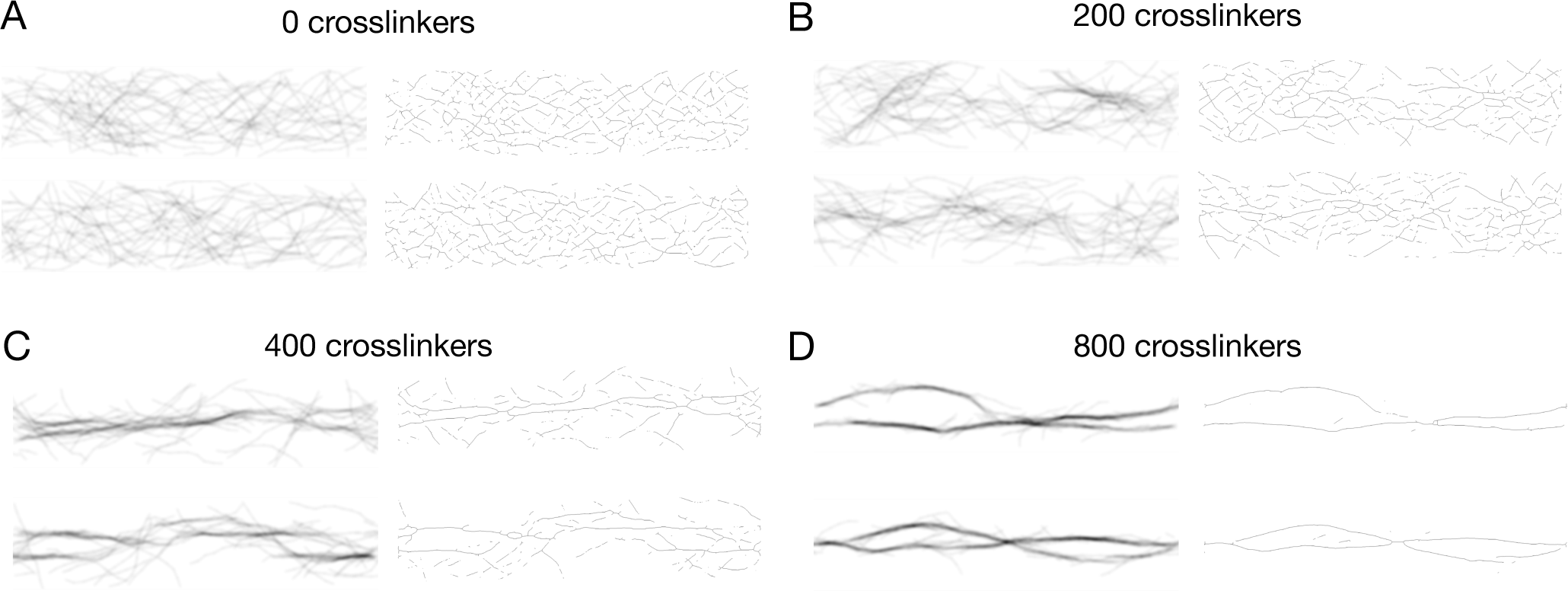
Snapshots of representative networks obtained from simulations. Simulated networks are shown for 0, 200, 400, and 800 crosslinkers (A – D). For each case, two sample networks are shown after 400 s of simulation. Pseudo-fluorescence images are shown at left, and skeletonized images are shown at right. Pseudo-fluorescence images have been inverted so that filaments appear dark against a white background.

While the image processing procedure generally captures the features of filament structures, there are also clear limitations. For example, the skeletonization procedure does not capture all of the filaments because of the thresholding of pixel intensities. In addition, both single filaments and larger bundles are represented as single-pixel lines, which can underrepresent the thickness of a bundle. However, the pixel intensities along the skeletonized features serve as a proxy for the number of bundled filaments. Overall, the skeletonization procedure captures the vast majority of prominent features of the actin network in the original fluorescence image.

### Morphometric analysis of network organization in simulated networks

Using the skeletonized images obtained from simulated actin networks, we calculated a variety of morphometric parameters that characterize different aspects of network organization. The parameters can be broadly classified as measures of filament *Density*, *Orientation*, *Ordering*, and *Bundling* (Table 1). These parameters include both published and unpublished parameters. To assess how well the parameters characterize the actual network organization, we computed analogous “ground-truth” parameters based on the actin filament positions from the raw simulation data. For the remainder of this paper, we refer to parameters as *measured* when they are obtained from image-based analysis and as *ground truth* when obtained directly from simulation data, where we know the location and shape of all filaments exactly. Details about the definitions and procedures for calculating the parameters can be found in Materials and Methods.

**Table 1:** Morphometric parameters.

| Category | Published parameters (units) | New parameters (units) | Ground truth (units) |
| --- | --- | --- | --- |
| Density | Occupancy (Higaki et al., 2020) | Distance ($\mu\text{m}$) | Occupancy<br>Distance ($\mu\text{m}$) |
| Orientation | Mean Angle ($^\circ$) (Madison et al., 2015) | Mean Weighted Angle ($^\circ$) | Mean Angle ($^\circ$) |
| Ordering | Parallelness (Ueda et al., 2010) | Angular Variation ($^\circ$)<br>Order Parameter | Parallelness<br>Angular Variation ($^\circ$)<br>Order Parameter |
| Bundling | Skewness (Higaki et al., 2010)<br>Coefficient of Variation (Higaki et al., 2020) | Bundle Parameter | Local Filament Bundling |

### Analysis of network *Density*

Measures of network *Density* provide information about the extent to which filaments occupy the domain containing the actin network. We calculated *occupancy* and *distance* for various numbers of crosslinkers. The *occupancy* is calculated as the fraction of pixels containing filaments within the skeletonized image (Higaki et al., 2010). The *distance* is the median distance from any pixel without a filament to the nearest filament.

Figures 2A and B show the measured and ground-truth values of *occupancy* respectively. Both the measured and ground-truth values decrease on average with increasing numbers of crosslinkers. To facilitate comparison between measured and ground-truth values, we plotted the measured values against ground-truth values for each simulated network (Fig. 2C). The relation between the measured and ground-truth values is approximately linear for *N_c_* ≤ 500, but it deviates at high levels of crosslinking. In this regime, the filaments are highly bundled and the measured values are less sensitive to changes in *N_c_* than the ground-truth values. The measured and ground-truth values of *distance* increase with more crosslinkers (Fig. 2D, Fig. S1), with a tight correlation of the values up to around *N_c_* = 600. The opposite trends for *occupancy* and *distance* arise because crosslinking promotes filament bundling, thereby consolidating filaments into bundles and leading to larger filament-free regions (Fig. 1). Although they follow the same trend, the measured *occupancy* values are 5 to 10 times smaller than ground-truth *occupancy* values. These differences can be attributed to the skeletonization procedure: Bundles that are several filaments thick are reduced to a single line, and thresholding leads to some filaments being neglected. The values of the measured and ground-truth *distance* are similar, indicating that the measured values are less sensitive to the image processing procedure.

**Figure 2.**
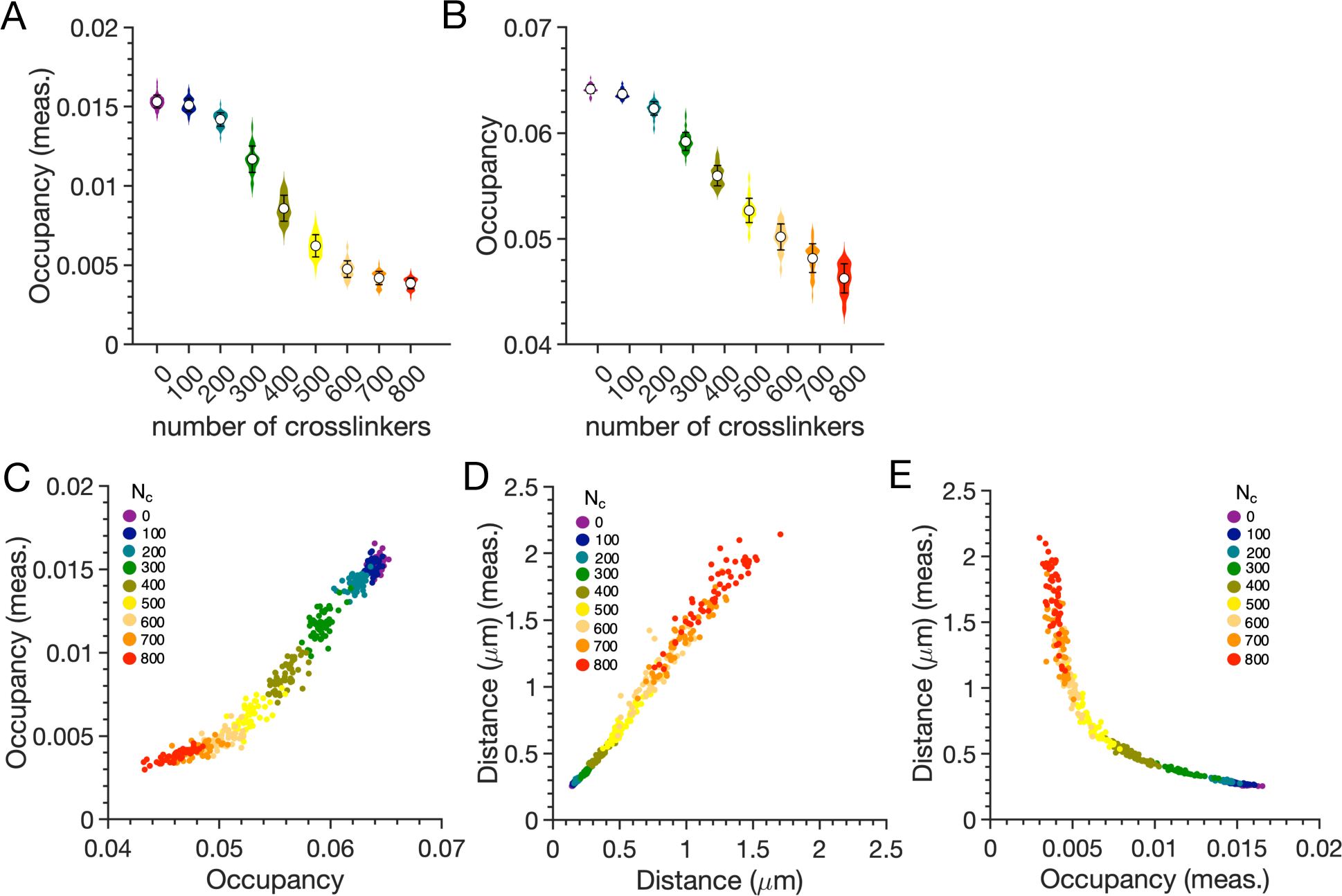
Distributions of the (A) measured and (B) ground-truth values of *occupancy* shown as violin plots for different numbers of crosslinkers. Means and standard deviations are shown. Each plot is constructed using 50 independent simulations for each value of *Nc*. (C, D) Network-by-network comparison of measured versus ground-truth parameters for network *Density*: (C) *occupancy* and (D) *distance*. Each point corresponds to a simulated network. (E) Network-by-network comparison of measured values of *distance* and *occupancy*.

We further compared the measured *occupancy* and *distance* values (Fig. 2E). This highlights that *occupancy* is much more sensitive than *distance* to changes in *N_c_* for more mesh-like networks (*N_c_* < 500). In contrast, *distance* is much more sensitive to changes in *N_c_* for highly bundled networks (*N_c_* > 500). Overall, the agreement between trends for measured and ground-truth parameters related to filament *Density* indicates that the measured parameters reliably capture information about the actual actin network, although this fidelity is reduced for highly bundled networks.

### Analysis of network *Orientation* and *Ordering*

Measures of network *Orientation* and *Ordering* provide information about the average orientation of filaments and the degree to which they are aligned with one another, respectively. We characterized network *Orientation* using the *mean weighted angle* of filaments relative to the long axis of the confinement area (Fig. S2). We determined the local angles by smoothing filaments over 0.5 µm segments (Madison et al., 2015), which reduced the variability of measurements based on pixel pairs that was used in a previously published algorithm (Ueda et al., 2010). In addition, this new method weighted the angle measurements by the pixel intensity to better reflect the number of individual filaments that have each measured orientation. These changes resulted in a new *Orientation* measure that better represented the ground truth data (compare Figs. S2C and D). Both measured values of *mean weighted angle* and ground-truth values of the *mean angle* remained close to zero regardless of crosslinker number. For bundled networks, this is a consequence of filaments preferentially aligning with the longitudinal axis of the simulation domain to minimize unfavorable bending (Akenuwa & Abel, 2023; Alvarado et al., 2014). For mesh-like networks at low crosslinker numbers, the mean angle near zero is simply a consequence of the random orientation of filaments.

To characterize the *Ordering* of filaments, we examined *angular variation*, *order parameter*, and *parallelness*. These parameters measure, respectively, the standard deviation of filament angles, the degree to which actin filaments align with the mean angle, and the degree to which actin filaments are parallel to each other. Figure 3 compares the measured and ground-truth values of these parameters for different numbers of crosslinkers. In contrast with the *mean weighted angle*, the *angular variation* decreased with increasing numbers of crosslinkers in both measured and ground-truth calculations (Fig. 3A, Figs. S3A and D). Furthermore, the *order parameter* (Fig. 3B, Figs. S3B and E) and *parallelness* (Fig. 3C, Figs. S3C and F) increased with increasing number of crosslinkers. These results indicate that with additional crosslinkers, filaments become more aligned, which is a consequence of the formation of bundles. Interestingly, we observed a kink in the shapes of *Ordering* parameter plots which gave rise to two approximately linear regimes. For both *angular variation* and *order parameter*, the kink occurred around *N_c_* = 300. For *parallelness*, the kink occurred around *N_c_* = 400. The kinks in the *order parameter* and *angular variation* are less pronounced than in *parallelness*. In addition, the *parallelness* measurements deviated more from the ground-truth data. This indicates that the *order parameter* and *angular variation* are better measures of ground-truth network *Ordering* than *parallelness*. For this reason, we exclude *parallelness* from our analysis below. In general, we observed good agreement between ground-truth and measured parameters of ordering.

**Figure 3.**
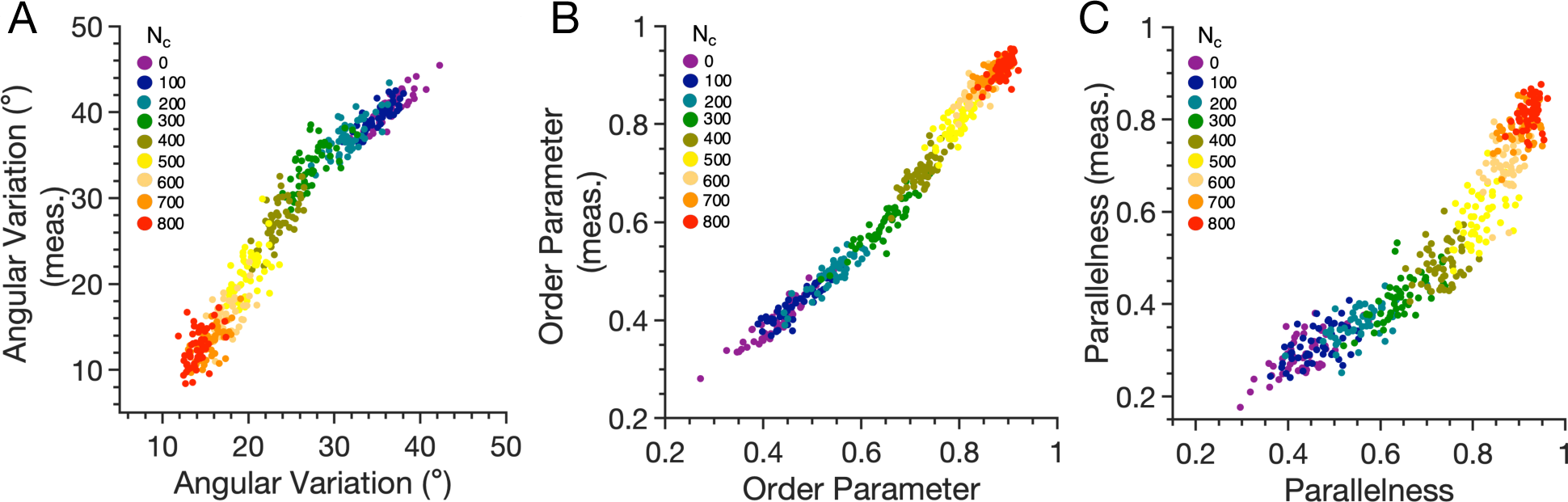
Network-by-network comparison of measured versus ground-truth parameters for network *Ordering*: (A) *Angular variation*. (B) *Order parameter*. (C) *Parallelness*.

### Analysis of network *Bundling*

Measures of network *Bundling* provide information about the degree to which filaments are organized into bundles. We first used two established metrics, the *skewness* and *coefficient of variation (CV)* of pixel intensities along the skeletonized filaments (Higaki et al., 2010; Higaki et al., 2020; Park & Nebenführ, 2013; Rosero et al., 2013; van der Honing et al., 2012). *Skewness* has been used widely as a measure of bundling, while *CV* was introduced more recently as an improved measure of bundling. To compare against a ground-truth value, we defined a metric termed *local filament bunding (LFB)*, which gives a measure of the characteristic bundle size as determined by the average number of filaments colocalized within small, occupied regions of the simulation domain. *LFB* increases with larger numbers of crosslinkers, highlighting that more filaments are within close proximity of one another in bundles (Fig. 4A). The distributions for large numbers of crosslinkers exhibit substantial overlap because adding crosslinkers to an already highly bundled system has only a marginal effect on the filament bundles.

**Figure 4.**
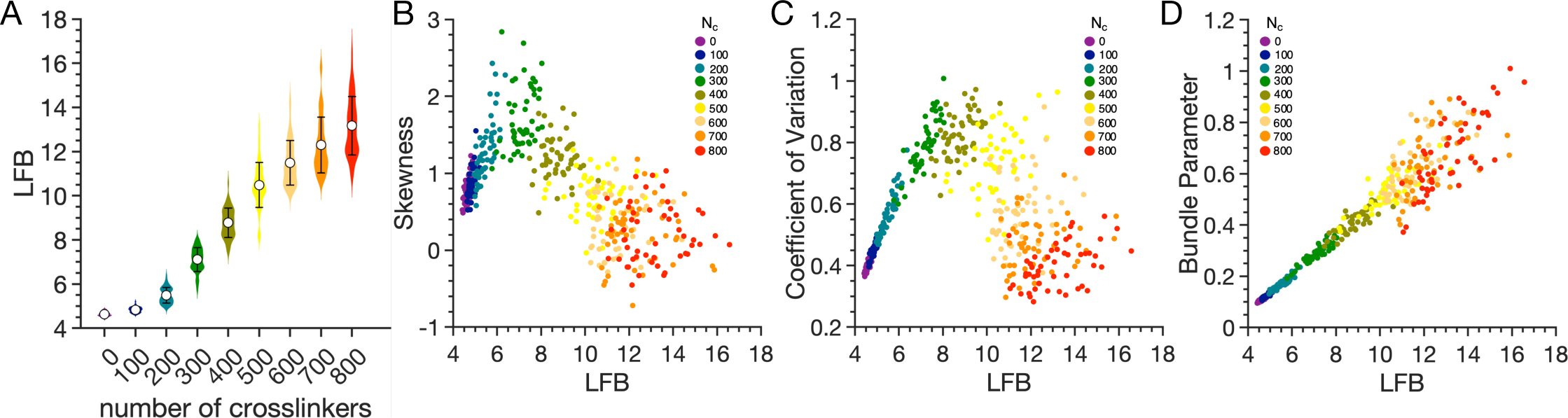
(A) Distributions of *local filament bundling* (*LFB*) shown as violin plots for different numbers of crosslinkers. Means and standard deviations are shown. (B-D) Network-by-network comparison of measured and ground-truth parameters for network *Bundling*: (B) *Skewness* of skeletonized filament pixel intensities. (C) *Coefficient of variation* of skeletonized filament pixel intensities. (D) *Bundle parameter*, defined as *coefficient of variation* multiplied by *distance*.

In contrast to *LFB*, both *skewness* and *CV* exhibit nonmonotonic behavior as a function of the number of crosslinkers (Figs. S4A and B). Plotting *skewness* and *CV* against *LFB* reveals that both suffer shortcomings in distinguishing the degree of bundling (Figs. 4B and C). The data indicate that *CV* is a better metric over a larger range, with a clear linear trend for smaller values of *N_c_*. This is consistent with reports suggesting that skewness is an unreliable measure of bundling, and that *CV* is a more robust measure in the cases where *skewness* breaks down (Higaki et al., 2020). However, our analysis indicates that *CV* also breaks down, following a non-monotonic trend with a peak around 400 crosslinkers (Fig. 4C, Fig. S4B). Thus, *CV* works well as a measure of bundling for networks with relatively low degrees of bundling, but it is not suitable as the degree of bundling increases.

Given that *skewness* and *CV* fail at high bundling levels, we sought to identify a measure that more closely reported the increase in *LFB* with increasing numbers of crosslinkers. Based on the drop of *CV* values at high levels of bundling and the corresponding increase in *distance*, we reasoned that multiplying *CV* by *distance* might compensate for the decrease of *CV* and provide better discrimination between cases. Thus, we defined a new parameter, the *bundle parameter*, as the *coefficient of variation* of skeletonized filament pixel intensities multiplied by *distance* (*CV* × *distance*). *CV* was chosen instead of *skewness* because it performed better as an indicator of bundling over a wider range of crosslinkers and is strictly nonnegative. In contrast to *skewness* and *CV*, *bundle parameter* values increased from 0 to 700 crosslinkers and maintained a strong linear correlation with ground-truth *LFB* before plateauing at very high numbers of crosslinkers (Fig. 4D, Fig. S4C). This result demonstrates that scaling CV by a measure of network *Density* provides a more robust measure of bundling, especially in highly bundled networks.

### Principal component analysis distinguishes between levels of crosslinking and reveals correlations in morphometric parameters

Morphometric parameters are useful in part because they can facilitate the grouping of images into different classes that reveal insight into the biology of the system being studied (Alderfer et al., 2022; Alizadeh et al., 2019; Pincus & Theriot, 2007). One approach is to use principal component analysis (PCA), which is a linear dimensionality reduction method. PCA can be used to visualize relationships between images based on their morphometric parameters, reveal correlations between the parameters, and help to identify distinct classes of networks.

We first performed PCA for the image-based parameters using 6 measured morphometric features for all 450 simulated networks: *occupancy*, *distance*, *mean weighted angle*, *angular variation*, *order parameter*, and *bundle parameter*. Figure 5A shows the data projected onto the first and second principal components. The first principal component (PC1) captured about 80% of the variance and yielded good separation between images associated with different levels of crosslinking. Networks with low levels of crosslinking were associated with negative values of PC1, while highly crosslinked networks were associated with positive values of PC1. The points associated with different levels of crosslinking were generally well separated except for the overlap of networks with small (0 and 100) and large (700 and 800) numbers of crosslinkers. This overlap reflects the inherent similarity between networks in these limits: With small numbers of crosslinkers, not enough filaments were bundled to yield significant differences in the morphometric parameters. With large numbers of crosslinkers, the vast majority of the filaments were bundled, leading to relatively minor differences as more crosslinkers were added.

**Figure 5.**
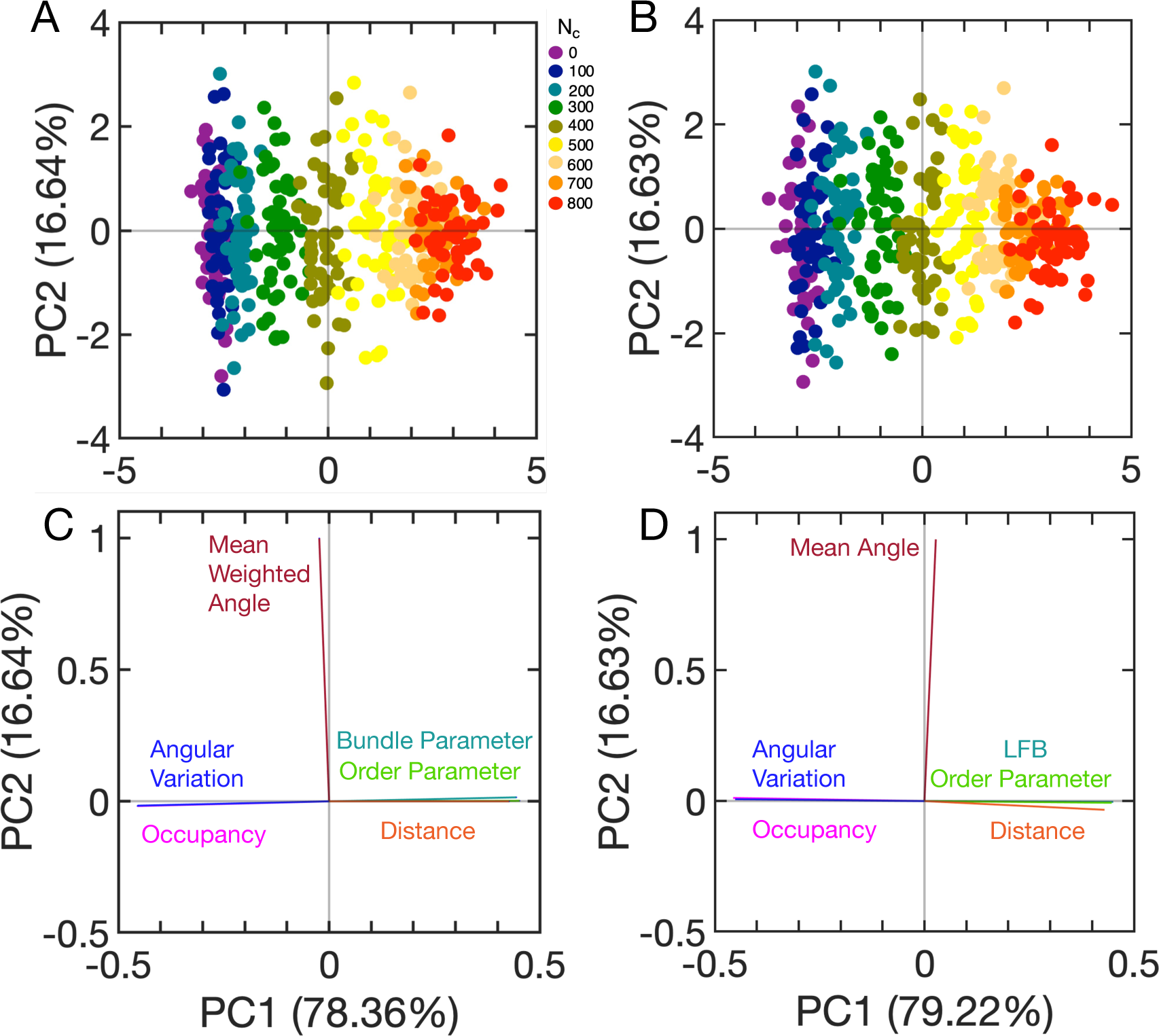
Principal component analysis of morphometric parameters obtained from simulated actin networks. (A, B) Projection of data onto the first two principal components (PC1 and PC2) using (A) measured morphometric parameters and (B) analogous ground-truth parameters. (C) Loading vectors for each measured morphometric parameter. (D) Loading vectors for each ground-truth parameter. The percent of variance explained by each principal component is reported in parentheses.

Measures of *Density*, *Ordering*, and *Bundling* contributed strongly to PC1, as indicated by their loadings (Fig. 5C). *Occupancy* and *angular variation* were highly correlated, and PC1 had negative loadings for these parameters. Hence, when the parameters increased, PC1 decreased. *Distance*, *bundle parameter*, and *order parameter* were highly correlated with one another but anticorrelated with *occupancy* and *angular variation*. PC1 had large positive loadings for these parameters, so when they increased, PC1 increased. Thus, increasing values of PC1 were associated with more bundled networks, which is consistent with larger numbers of crosslinkers being associated with larger values of PC1 (Fig. 5A). The correlations between morphometric parameters were also reflected by pairwise correlations coefficients (Table S1). PC2 was dominated by the *Orientation* parameter of *mean weighted angle*. However, PC2 captured a relatively small fraction of the overall variance and showed no obvious discrimination between the various degrees of crosslinking. Thus, for the networks considered here, this measure was not useful for categorizing networks. We speculate that the second principal component primarily reflects the inherent variability between networks even at the same conditions.

For comparison, we also performed PCA using the analogous ground-truth morphometric parameters (Fig. 5B). The results were similar, reflecting the high degree of correlation seen in the previous comparison of measured and ground-truth parameters. As before, the first principal component captured about 80% of the variance and yielded similar separation between networks with different numbers of crosslinkers. The loading plot revealed that analogous ground-truth parameters similarly contributed to PC1 and PC2 (Fig. 5D). Taken together, the measured and ground-truth PCA results show that the morphometric parameters quantitatively characterize aspects of actin network organization that can be used to distinguish between different classes of networks that differ in their degree of bundling.

### Application of morphometric parameters to natural actin networks in root epidermal cells

Having validated a useful set of morphometric parameters with synthetic actin networks, we tested whether the parameters allowed us to distinguish different actin networks in root epidermal cells. For this purpose, we imaged the outer cortical actin network in fully expanded atrichoblasts in the early differentiation zone of the root epidermis in young *Arabidopsis thaliana* seedlings in both wild-type and *myo11e* myosin mutants (Park & Nebenführ, 2013). Analysis of the confocal images followed the same procedure described above for the pseudo-fluorescence images based on our synthetic networks. To further simplify the analysis of the images, we introduced an automated detection of the cell area based on the convex hull of all detected actin filaments. This ensured that the calculation of *Density* measures was based on the actual visible cell area and not on the arbitrary size of the image frame.

The actin networks showed considerable variation between cells (Fig. 6). In both genotypes, we observed cells with a dense mesh of actin filaments as well as cells with a less dense network. All cells also contained a combination of thin and faint filaments, and wider and brighter filaments that presumably represent bundles containing multiple actin filaments. The complete set of 30 cell images per genotype used for the quantitative analysis is shown in Figure S5.

**Figure 6.**
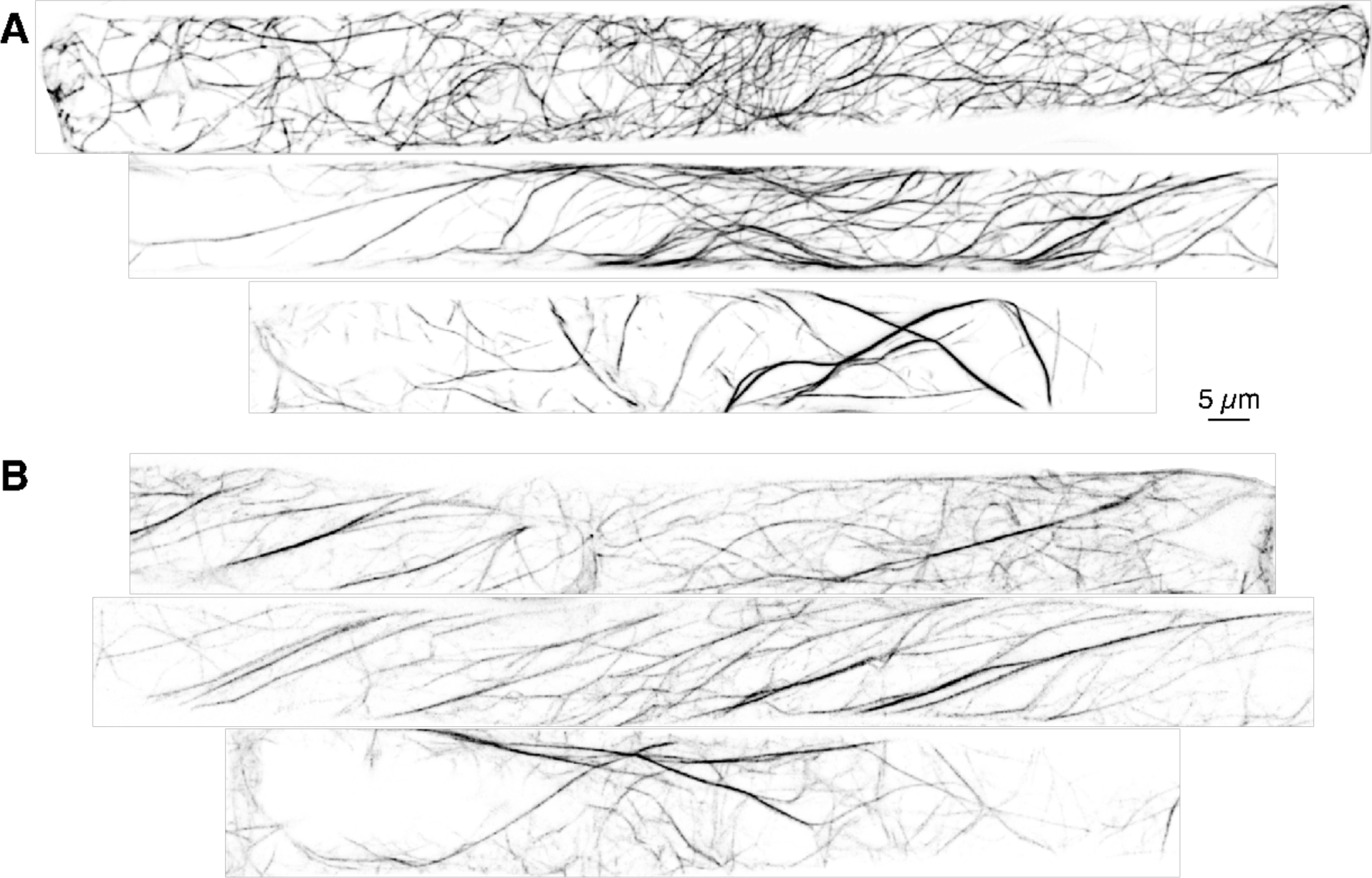
Representative images of actin networks in root epidermis cells of (A) wild-type and (B) myosin mutant seedlings. Fluorescent images have been inverted so that filaments appear dark on a bright background.

Comparison of morphometric parameters obtained from images of wild-type and *myo11e* mutant roots revealed significant reduction in *occupancy* in the mutant cells (Fig. 7A). Unexpectedly, the other *Density* measurement, *distance*, did not show a significant difference between the genotypes (Fig. 7B). The *Orientation* of the filaments (*mean weighted angle*) was not different between the genotypes (Fig. 7C). The two *Ordering* parameters, *angular variation* and *order parameter* (Figs. 7D and E), indicated that the mutant networks were significantly more ordered than those found in wild-type cells. Similarly, the *bundle parameter* was on average higher in the *myo11e* cells compared to wild-type cells (Fig. 7F).

**Figure 7.**
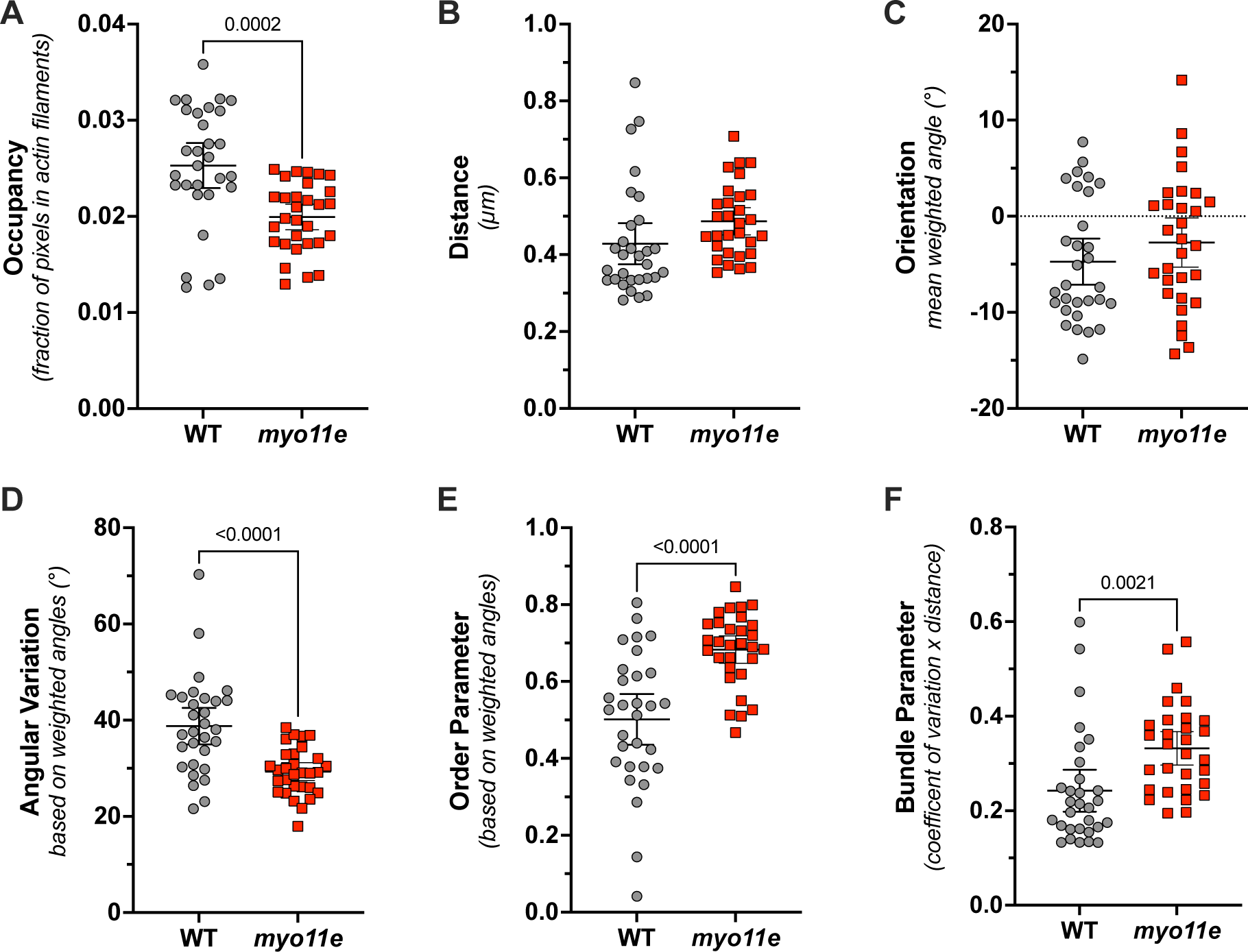
Image analysis of actin networks in root epidermis cells of wild-type (WT) and myosin mutant (*myo11e*) seedlings. (A) *Occupancy*. (B) *Distance*. (C) *Orientation*, measured as *mean weighted angle*. (D) *Angular variation*. (E) *Order parameter*. (F) *Bundle parameter*, measured as the *coefficient of variation* of skeletonized filament pixel intensities multiplied by *distance*. Welch’s *t*-test was used to calculate *p*-values. Statistically significant results are shown.

To further explore the relationship between genotypes and morphometric parameters, we performed PCA using these six parameters from the 60 experimental images (Fig. 8). Although there is overlap between wild-type and *myo11e* mutant cells in the PC1-PC2 plane, there is clear clustering of each cell type (Fig. 8A). Both PC1 and PC2 had substantive contributions from *Density*, *Ordering*, and *Bundling* parameters, and together they captured about 80% of the variance (Fig. 8B). In contrast, for the simulated networks, only PC1 had substantial contributions from these parameters. Interestingly, PC1 for the experimental images had loadings similar to those for PC1 from the simulations, while PC2 mainly had large contributions from *Ordering* parameters. This suggests that PC1 captures features of the networks similar to those in the simulated networks, while PC2 captures distinct features of the ordering of filaments in the actin networks. PC2 does not have an analogous principal component in the simulation data, which suggests that there are meaningful features in the experimental images that are not present in the simulation data.

**Figure 8.**
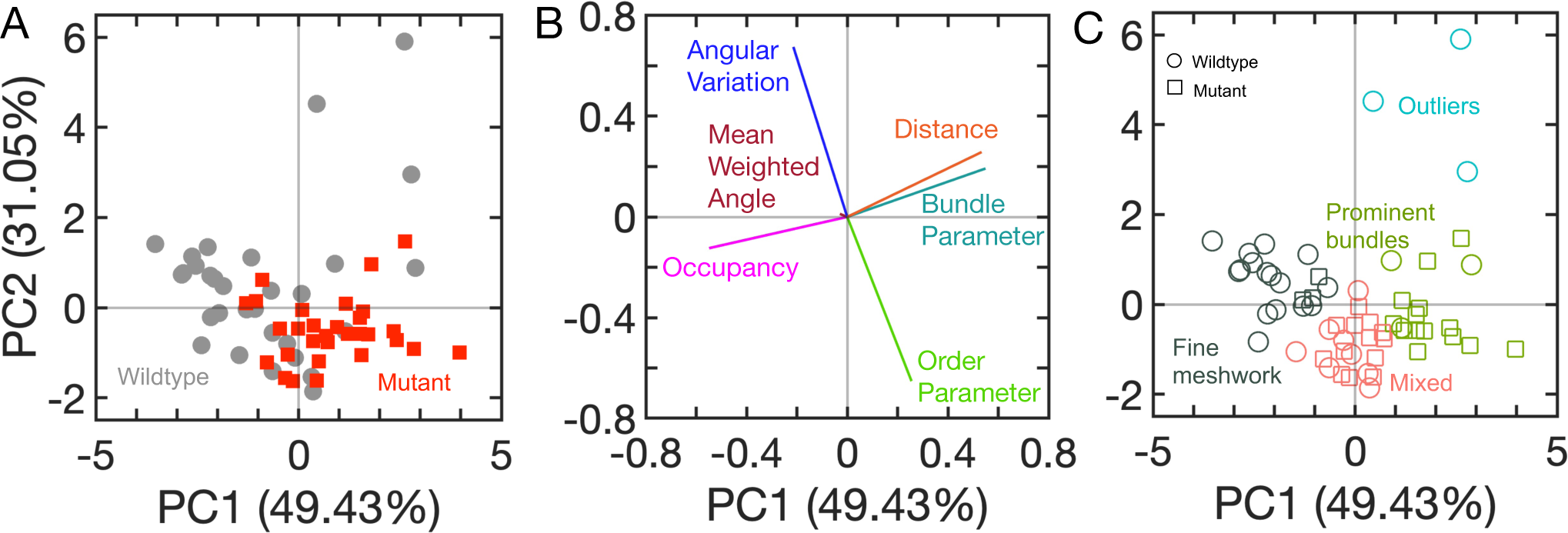
Principal component analysis of morphometric parameters derived from actin networks in root epidermis cells of wild-type (WT) and myosin mutant (*myo11e*) seedlings. (A) Data from cells projected onto the first two principal components (PC1 and PC2). Wild-type cells are shown as grey circles. Mutant cells are shown as red squares. The percent of variance explained by each principal component is reported in parentheses. (B) Loading vectors for each morphometric parameter. (C) Result of k-means clustering (k=4) on the projected data with each cluster shown in a different color. Wild-type cells are shown as circles and mutant cells are shown as squares. The fine-meshwork cluster (dark green) contains 16 wild-type and 3 mutant cells, the mixed cluster (orange) contains 8 wild-type and 13 mutant cells, the prominent-bundles cluster (light green) contains 3 wild-type and 14 mutant cells, and the outlier cluster (blue) contains 3 wild-type cells.

Interestingly, even though *Bundling*, *Density,* and *Ordering* parameters all indicated clear differences between wild-type and mutant cells, there was a striking lack of correlation between the *Ordering* parameters and the other two categories (Table S2). In the PCA, this was reflected by the nearly perpendicular loading vectors for *Ordering* parameters on one hand, and *Density* and *Bundling* on the other (Fig. 8B). *Mean weighted angle* was found to be largely independent of all other measurements and was the dominant contributor to PC3 in the PCA analysis. This is analogous to its contribution to PC2 in the simulated networks, and it again does not appear to be an important factor in distinguishing between different classes of networks.

We used k-means clustering with k = 4 to cluster the data points in the PC1-PC2 plane. This resulted in three primary clusters distinguished by different levels of fine meshwork and local bundles. A fourth cluster (“outliers”) corresponded to large values of PC2 and was dominated by very prominent bundles and larger filament-free regions (Figs. 8C and S6D). Of the primary clusters, the cluster with a dense network of fine actin filaments and a limited number of brighter bundles (“fine meshwork”) was comprised mainly of wild-type cells (Fig. S6A). A second “mixed” cluster, which contained a mixture of both wild-type and *myo11e* mutant cells, was characterized by more prominent filament bundles together with areas of dense filament meshwork (Fig. S6B). A third cluster, characterized by bright bundles and a less dense meshwork of fine filaments, was comprised largely of *myo11e* mutant cells (Fig. S6C). The three primary clusters were sorted largely along PC1, with large values corresponding to more bundled networks. Importantly, both genotypes were found in all clusters, suggesting that both genotypes were able to generate networks with different architectures.

## Discussion

We set out to test the validity and usefulness of morphometric parameters that can be extracted from images of fluorescently labeled actin filaments. Using computer simulations, we generated actin networks that have a striking resemblance to those of natural cells. With these networks, we constructed pseudo-fluorescence images analogous to those from experiments. In contrast with previous studies utilizing morphometric parameters, our simulation results allowed us to test the fidelity of the parameters by comparing them to ground-truth parameters determined from the underlying network of filaments.

A crucial aspect of analyzing fluorescence images of actin networks is the identification of actin filaments. Simple thresholding based on image brightness works only for ideal images with minimal noise and uniform background, and it typically leads to problems with the detection of filaments in denser networks. We have implemented an approach that uses multi-directional linear convolutions to emphasize linear components of the filament network to allow for more robust detection of faint filaments by thresholding (Sun & Vallotton, 2009). This approach still does not capture all actin filaments, but it is able to detect both bright bundles and faint signals. Our implementation of this approach in an ImageJ macro is similar to that described by Higaki and coworkers (Higaki et al., 2010, 2020). Although the two algorithms result in subtle differences in the skeletonized images, the disparities lead only to small differences in measured parameters (data not shown).

We categorized the morphometric parameters into four main categories: *Density*, *Orientation*, *Ordering*, and *Bundling*. In general, the measured values derived from pseudo-fluorescence images were highly predictive of the analogous ground-truth parameters, although the values were not always consistent, and the relation was not always linear. For *Orientation* and *Ordering* parameters, the measured parameters were close in value to the ground-truth parameters. The measures of filament *Density*, on the other hand, showed strong correlation with the ground-truth data but underestimated the actual extent of the network: measured *occupancy* values were lower than ground-truth values and *distance* values where larger than ground truth. For *Bundling* parameters, we compared measured values to *local filament bundling*, which characterized the average bundle size in the ground-truth networks. Existing methods suffered shortcomings, so we introduced a new *bundle parameter* that provided a useful measure of filament bundling across all crosslinking levels. All morphometric parameters measured here, irrespective of the differences in absolute values between measured and ground-truth values, generally provide a good representation of the ground truth.

The comparison to ground truth data allowed us to detect shortcomings in some of the published morphometric parameters. For example, the *mean angle* calculation described in Ueda et al. (Ueda et al., 2010) systematically overestimated deviations from the horizontal and contained more noise compared to the angle calculation based on a sliding window employed here (Fig. S2D). The calculation of local angles along the filaments enabled us to calculate more accurate measures for network *Ordering*, namely *angular variation* and *order parameter*. These parameters matched the ground truth values more closely than *parallelness*, which is based on frequencies of pixel pairs and therefore is inherently more noisy (Fig. 3). In addition, *parallelness* consistently underestimated the ordering of the networks. The largest discrepancy with ground truth values was found for the *Bundling* measures of *skewness* and *coefficient of variation*. Both measures displayed a nonmonotonic relation with *local filament bundling* and decreased at higher levels of crosslinking. Thus, for a given parameter value, it would be impossible to identify the degree of bundling based on the value of these *Bundling* parameters alone. While *skewness* and particularly *coefficient of variation* remain useful *Bundling* measures for networks dominated by fine meshwork, we recommend using *bundle parameter* since it provides a reliable estimate of bundling levels over a much broader range.

The differences between measured and ground-truth values of *occupancy* and *distance* were likely caused by the image processing pipeline, where thresholding eliminated faint signals from single filaments and skeletonization underrepresented the true width of actin bundles. Further, *occupancy* was less sensitive to the level of crosslinking in highly crosslinked networks compared to less crosslinked networks. In contrast, *distance* remained sensitive to changes in the number of crosslinkers in highly crosslinked networks. Plotting measured *distance* against measured *occupancy* (Fig. 2E) revealed that each parameter was most sensitive to changes in the level of crosslinking in different regimes, thus suggesting that there is value to calculating both when analyzing images of filament networks.

Curiously, several of the measured parameters showed a deviation from a strictly linear response when plotted against ground-truth values. Although the deviation was most pronounced for *occupancy* (Fig. 2A), it can also be seen for *angular variation* (Fig. 3A) and to some extent for *order parameter* (Fig. 3B). This may be an effect of the skeletonization procedure that misses some aspects of the networks depending on their density or bundling level which prevents accurate representation of the filaments in the single-pixel lines.

When applying the morphometric measures to natural actin networks, it is important to bear in mind certain limitations. In the simulations, the number of filaments, and hence the total fluorescence signal, was constant between the different conditions. However, we cannot know whether a similar situation exists in plant cells of different genotypes or under different treatment conditions. This problem can, at least partially, be avoided by enhancing linear features prior to applying a threshold based on the signal intensities present in the individual image as implemented here. Nevertheless, variable signal intensity between samples can make comparisons difficult since it is not known whether single filaments can be reliably detected. In general, it is therefore advisable to keep exposure settings (e.g., laser power and detector sensitivity) constant between images as long as saturation of pixels can be avoided since this will affect the calculation of the *bundle parameter* which depends on proper determination of signal variation.

Interestingly, most measured parameters of natural networks fall into the range of zero to 500 crosslinkers in our simulated networks, suggesting that the natural networks in the observed cells do not reach the very high bundling levels we imposed in our simulations. The only deviation from this general rule was the *occupancy* values, which were slightly higher in biological networks. This was likely caused in part because the reference area is limited to the convex hull around the detected filaments in experimental images. Filament anchoring in cells, which is not a feature of the simulations, may also play a role. This may be a useful feature to include in future simulations of actin networks.

The analysis of biological data revealed clear differences between the actin networks in wild-type and *myo11e* mutant cells (Fig. 7). Curiously, our results suggest that the myosin mutant has more bundling in its root epidermal cells than wild-type, which is different from findings using double mutants (Madison et al., 2015; Ueda et al., 2010). This unexpected effect of the myosin mutation might reflect inherent differences between the cell types observed in the different studies. Alternatively, the level of bundling may be different for single and double mutants. Importantly, none of the morphometric parameters by itself provided a simple, clear-cut distinction between the genotypes. This overlapping distribution of the two genotypes was also found in the principal component analysis, suggesting that the actin filament networks of the two genotypes are not fundamentally different, but that the mutation simply shifts the balance more towards bundling (Fig. 8). These differences in actin organization did not appear to affect root elongation of the seedlings as both genotypes showed similar growth rates.

Another critical result from principal component analysis is the discovery that *Ordering* parameters are only weakly correlated with *Density* and *Bundling* parameters, in contrast with the simulation data. This suggests that cells can regulate filament organization independently of bundling levels, which likely requires additional regulatory elements that we had not included in our simulations. One candidate for such a regulatory factor could be proteins that anchor actin filaments to other elements in a cell, for example, the nucleus or the plasma membrane. It is also conceivable that the three-dimensional nature of the cytoplasm, which forms a continuous sleeve around the central vacuole, leads to other actin network configurations than can be generated in the 2D plane used in our simulations. The number and length of actin filaments was also fixed in simulations, which is in contrast with the dynamic nature of filaments in cells.

Both the synthetic networks and the natural networks analyzed here had filaments that were on average aligned with the long axis of the confinement area or cell, respectively. As a result, the *Orientation* parameter was not useful to distinguish the different networks. However, it is conceivable that different genotypes or treatments may lead to a predominant orientation that deviates from the long axis of the cell. For example, loss of multiple myosin XI genes has been found to lead to a nearly complete loss of longitudinal actin bundles in leaf midvein epidermal cells and the appearance of oblique or transverse bundles (Peremyslov et al., 2010). The *mean weighted angle* measure would be able to quantify these changes.

Taken together, our results indicate that image-derived morphometric parameters provide a useful measure of actin network organization that is faithful to the underlying organization of actin filaments. Further, by combining several measures, one can distinguish subtle differences between cells, thus emphasizing the importance of morphometric analysis in helping to decipher how cytoskeletal organization is influenced by cell identity, genotype, or other experimental perturbations.

## Materials and Methods

### Computer simulations of crosslinked actin networks

We utilized the Actin Filament Network Simulation (AFINES) model to simulate crosslinked actin filaments in confined environments (Freedman et al., 2017). Details of the simulations can be found elsewhere (Akenuwa & Abel, 2023; Freedman et al., 2017). Briefly, AFINES is a coarse-grained model that uses kinetic Monte Carlo and Brownian dynamics methods to simulate the dynamics of actin filaments and actin crosslinking proteins in two dimensions. Actin filaments are modeled as semiflexible, bead-spring polymers with a bending stiffness chosen to match the persistence length of actin filaments. Crosslinking proteins are modeled as springs with two ends that can stochastically bind and unbind from filaments. Crosslinkers preferentially crosslink filaments that are locally aligned with a small relative angle between them (Akenuwa & Abel, 2023). Dynamics are governed by alternating between a kinetic Monte Carlo step to update the binding and unbinding of crosslinkers and a Brownian dynamics step to update the positions of filaments and crosslinkers.

Actin filaments and crosslinkers were simulated in a 40 µm × 10 µm domain to reflect the characteristic shape of root epidermal cells. The simulations used reflective boundary conditions in the short dimension and periodic boundary conditions in the long dimension. We simulated a fixed number of actin filaments (100) of length 10 µm and varied the number of crosslinkers, with *N_c_* = 0, 100, 200, 300, 400, 500, 600, 700, and 800. We generated 50 independent samples for each simulation condition. Each network was simulated for 400 s, and the final configuration was used for analysis.

### Generation of pseudo-fluorescence images from simulations

To mimic confocal microscopy images, we converted simulated actin networks into pseudo-fluorescence images. To do this, we placed point sources every 0.1 µm along each filament to mimic fluorescent markers. The system was divided into 0.1 µm × 0.1 µm voxels, and the intensity of each source was modeled as a Gaussian function with σ = 0.1 µm. We added the intensity from each source for all voxels within 0.2 µm of the source, and the resulting intensity map was plotted as an image with a simulated pixel size of 0.0212 µm.

### Identification of actin filaments in pseudo-fluorescence and confocal images

Fluorescence images were processed similarly to Higaki et al. (2010). Briefly, the original images were smoothed with the “Mean…” filter in ImageJ and a smoothing radius of 0.2 µm. The resulting image was further processed by subtracting the background with a sliding paraboloid algorithm, followed by applying multidirectional non-maximum suppression to enhance the linear features of the image (Sun & Vallotton, 2009). This enhanced image was thresholded at the mean intensity of all non-zero pixels and converted to a binary image.

Subsequent skeletonization was performed using the “Skeletonize” command in ImageJ, yielding single-pixel lines of uniform intensity. This image was used as a mask to extract the pixel intensities from the original, unprocessed image.

### Determination of measured morphometric parameters from images

We calculated morphometric parameters from confocal images of root epidermal cells and pseudo-fluorescence images of simulated actin networks. We organized the morphometric parameters into four categories: network *Density*, *Orientation*, *Ordering*, and *Bundling*.

#### Measures of network *Density*

Network *Density* was measured using *occupancy* and *distance*. *Occupancy* was calculated as the fraction of pixels containing filaments within the skeletonized image (Higaki et al., 2020). *Distance* was calculated as the median distance from each pixel without a filament to the nearest filament. The distribution of distance values was obtained with the “Distance Map” command in ImageJ.

#### Measure of network *Orientation*

Network *Orientation* was captured by the mean of filament angles weighted by pixel intensities. To obtain the angles of filaments, we employed a sliding window approach. For every pixel in the skeletonized filaments, we moved along pixels comprising a continuous filament to cover a length of 0.5 µm. A line segment was drawn between the endpoints, and the angle relative to the horizontal was calculated (Madison et al., 2015). We obtained the *mean weighted angle* by multiplying each angle by the intensity (weight) of the central pixel and averaging with respect to the total weight. Weighting by the intensity of the pixels provides a measure of the number of actin filaments represented by the single skeletonized filament.

#### Measures of network *Ordering*

Network *Ordering* was measured using the standard deviation of filament angles (*angular variation*) and an *order parameter* based on skeletonized filament orientation. The *angular variation* was calculated as the standard deviation of the weighted distribution of angle measurements above. To obtain the *order parameter*, we utilized the same sliding window approach as above and calculated 〈2 cos^2^(*θ*) − 1〉, where *θ* is the angle of the line segment relative to the mean weighted angle and ⟨·⟩ denotes a weighted average over all pixels in the skeletonized filaments. To calculate *parallelness*, the angles of all neighboring pixel pairs in the skeletonized image were determined as 0, 45, 90, and 135° relative to the longitudinal axis of the image, as described in Ueda et al. (2010). The *parallelness* is given by

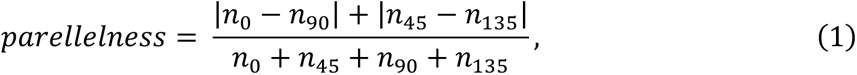

where *n*_0_, *n*_45_, *n*_90_, and *n*_135_ are the numbers of pixel pairs that make 0, 45, 90, and 135° angles, respectively.

#### Measures of network *Bundling*

*Bundling* of filaments was measured using the *skewness* (Higaki et al., 2010) and *coefficient of variation (CV)* (Higaki et al., 2020) of the distribution of pixel intensities within the skeletonized filaments. We also calculated a new *bundle parameter*, defined as *CV* × *distance*.

### Determination of ground-truth morphometric parameters from simulated actin networks

Analogous ground-truth morphometric parameters were calculated using the simulated actin filament configurations. We determined the *occupancy* of a sample by dividing the system into voxels of side length 0.0212 µm and calculating the fraction of voxels containing at least one filament.

To calculate *distance*, we determined the distribution of distances from all voxels without a filament to the closest voxel with a filament. *Distance* was the median value of the distribution.

To obtain the *mean angle* and *angular variation*, we determined the angle of each 1-µm segment along the filaments relative to the horizontal axis and then computed the mean and standard deviation of all such angles. The *order parameter* was calculated as 〈2 cos^2^(*θ*) − 1〉, where *θ* is defined as the angle relative to the mean angle and 〈·〉 denotes an average over all segments. The *parallelness* was calculated using equation (1) above with *n*_0_ the number of angles between 0 and 22.5° and 157.5 and 180°, *n*_45_ the number of angles between 22.5 and 67.5°, *n*_90_ the number of angles between 67.5 and 112.5°, and *n*_135_ is the number of angles between 112.5 and 157.5°.

To characterize network *Bundling*, we defined a parameter termed *local filament bundling (LFB)*. We divided the system into voxels of side length 0.025 µm and identified the filaments in each voxel. Then, for each voxel containing a filament, we placed a 0.5 µm × 0.5 µm box centered on the voxel and determined the number of unique filaments in that box. All such values were averaged to obtain the *LFB*. A larger value of *LFB* indicates that more filaments are within close proximity on average, thus indicating a higher degree of bundling.

### Code availability

Simulation and analysis code will be readily provided upon request.

### Experimental details

#### Plant material

All *Arabidopsis thaliana* seeds were obtained from the Arabidopsis Biological Resource Center (ABRC; abrc.org). Columbia (Col-0) was used as wild-type. Myosin XI mutants used were *myo11e-3* (*xik-3*; SALK_018764) (Ojangu et al., 2007) All plant lines were transformed with the YFP-FABD2 actin marker driven by the double 35S promoter (Park & Nebenführ, 2013). Seedlings were grown vertically on ¼ MS medium with 1% sucrose and 0.5 % phytagel for 5 days prior to imaging.

#### Confocal Imaging

Seedlings were placed in a 5 cm dish with a #1.5 cover glass bottom and covered with a small patch of phytagel in growth medium. Fully grown root epidermal cells were imaged on a Leica SP8X laser scanning confocal microscope with a 63x/1.4 oil objective. Imaging of the YFP fluorophore used the 514 nm line of the white-light laser and a HyD detector with a detection window from 520 nm to 590 nm. The outermost 4 to 7 µm of epidermal cells was imaged as a Z-stack with 0.2 µm step size and a pixel size of about 45 nm. The resulting image stack was deconvolved with the adaptive “Lightning” process in Leica LAS X software and collapsed into a single 2D image by maximal intensity projection.

## Acknowledgements

This work was supported by the National Science Foundation (MCB-1715794). We thank Jaydeep Kolape in the Advanced Microscopy and Imaging Center for technical assistance.

