## Supplemental figures for "Morphometric Analysis of Actin Networks"

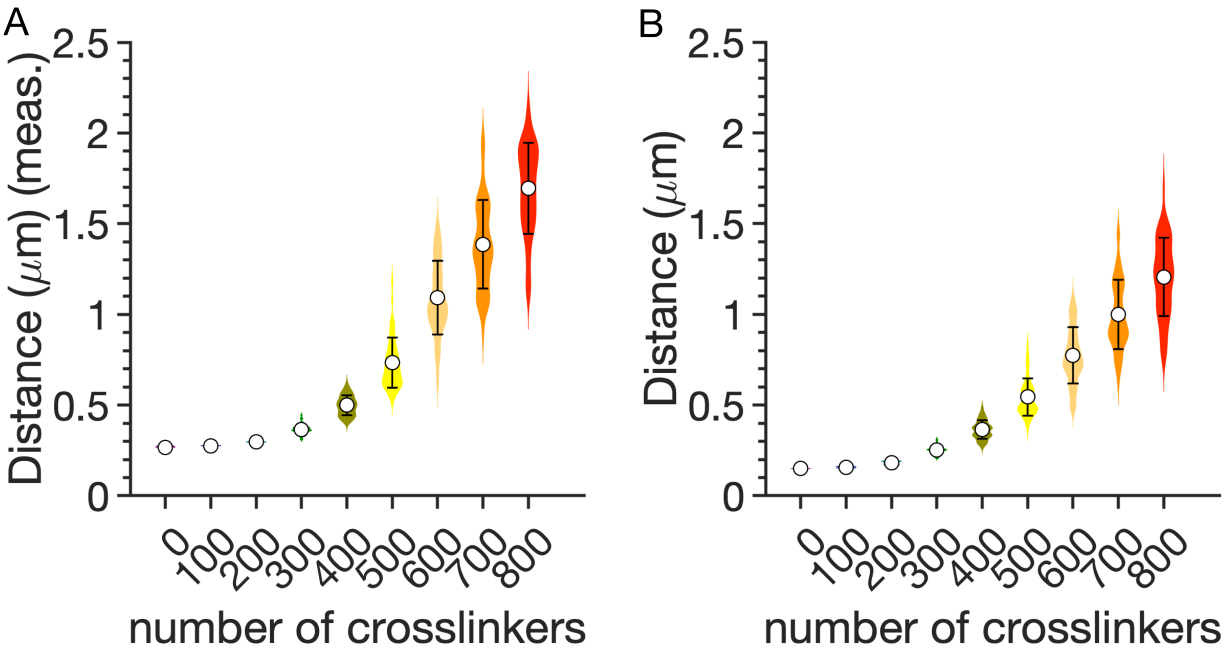


**Figure S1**. Distributions of (A) measured and (B) ground-truth *distance* shown as violin plots. Means and standard deviations are shown. Each plot is constructed using 50 independent simulations for each value of $N_c$.

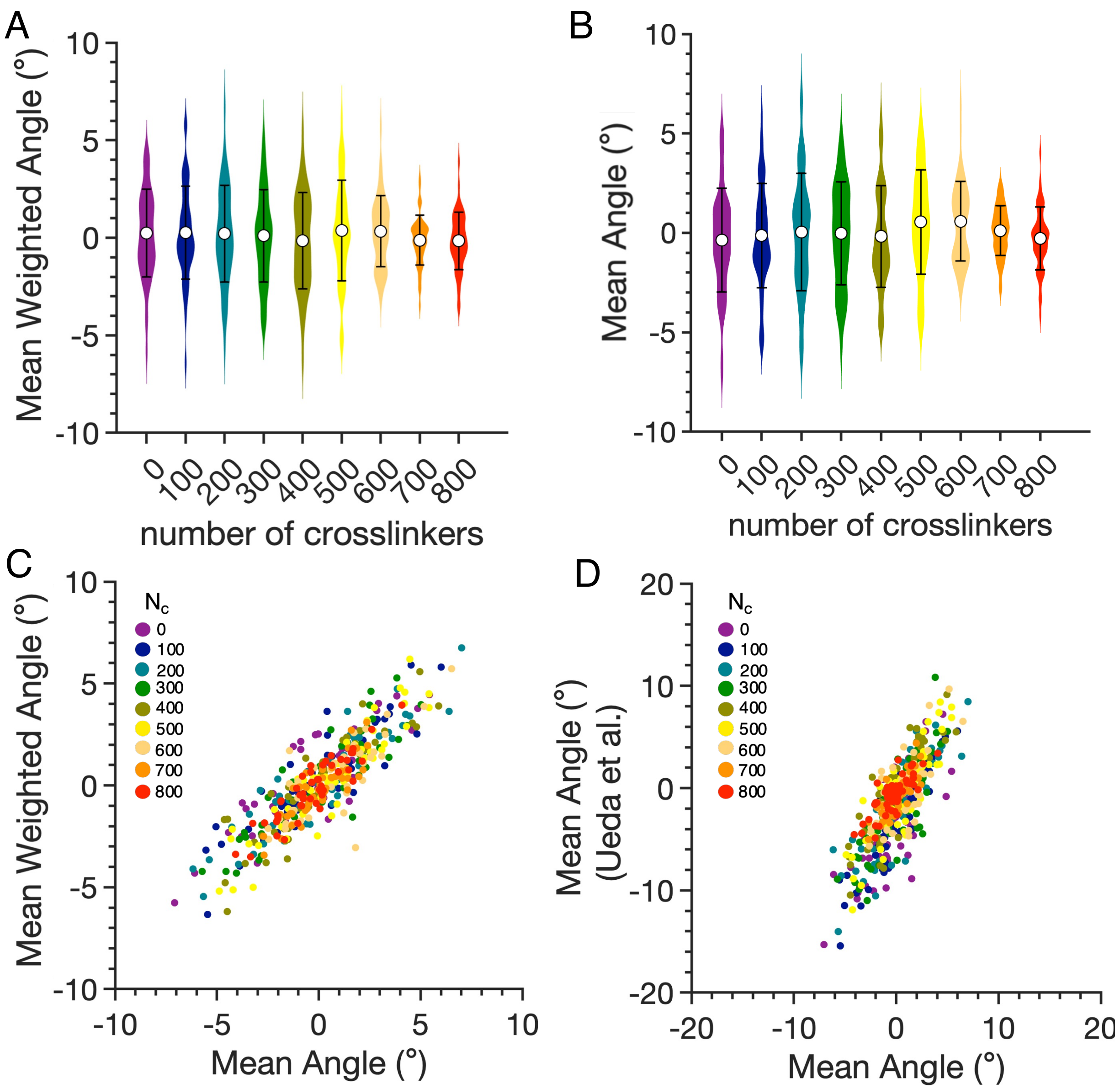


**Figure S2**. Distributions of (A) measured *mean weighted angle* and (B) ground-truth *mean angle* shown as violin plots. Means and standard deviations are shown. (C) Network-by-network comparison of measured *mean weighted angle* and ground-truth *mean angle*. (D) Network-by-network comparison of *mean angle* calculated using the method of Ueda et al. and ground-truth *mean angle*.

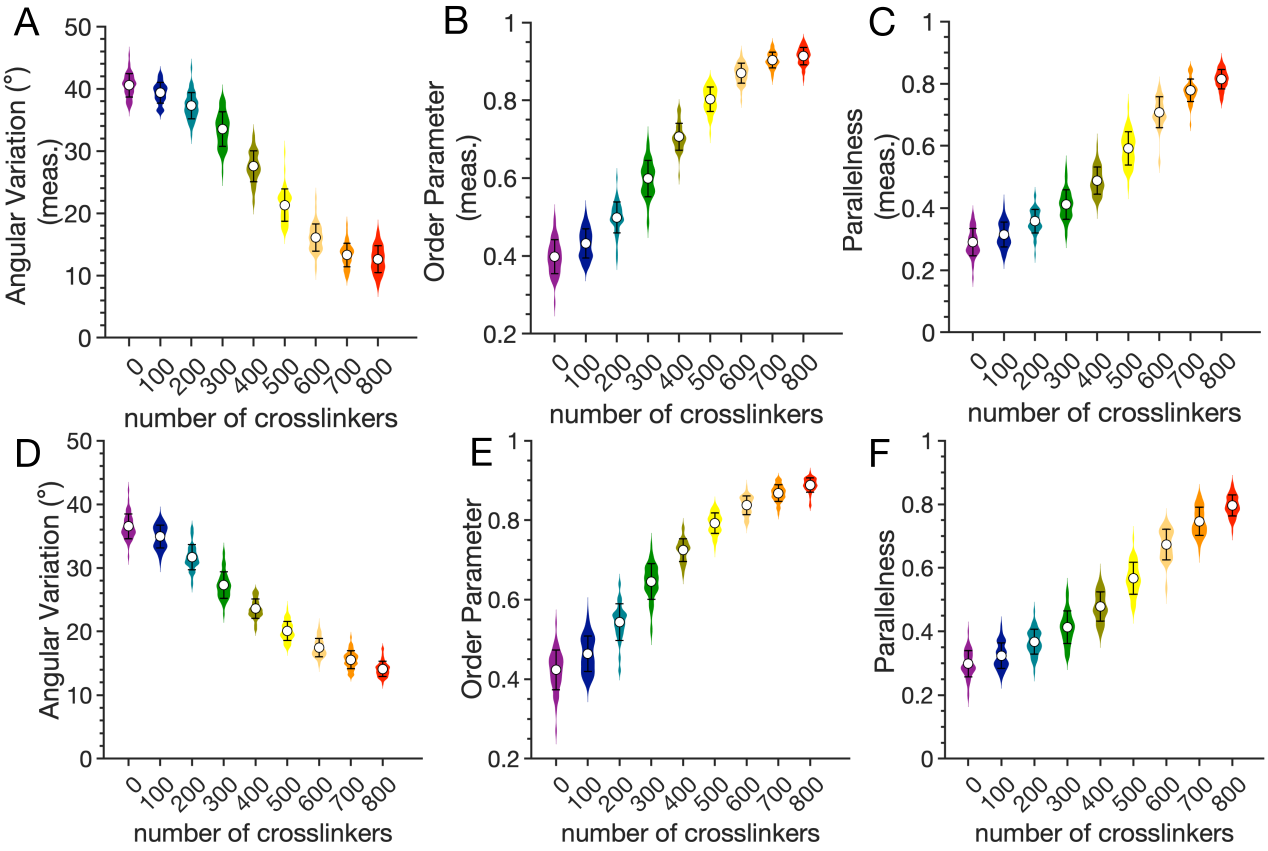


**Figure S3**. Distributions of measured and ground-truth *Ordering* parameters shown as violin plots. Means and standard deviations are shown. Top row: Measured parameters. (A) *Angular variation*. (B) *Order parameter*. (C) *Parallelness*. Bottom row: Ground-truth parameters. (D) *Angular variation*. (E) *Order parameter*. (F) *Parallelness*.

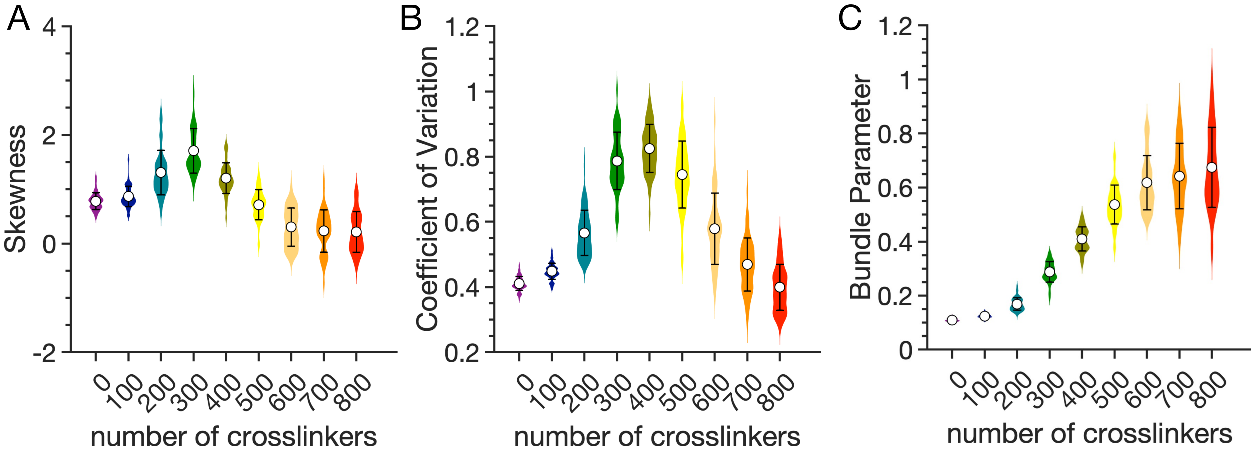


**Figure S4**. Distributions of measured *Bundling* parameters shown as violin plots. Means and standard deviations are shown. (A) *Skewness* of skeletonized filament pixel intensities. (B) *Coefficient of variation* of skeletonized filament pixel intensities. (C) *Bundle parameter,* defined as *coefficient of variation* of filament pixel intensities multiplied by *distance*.

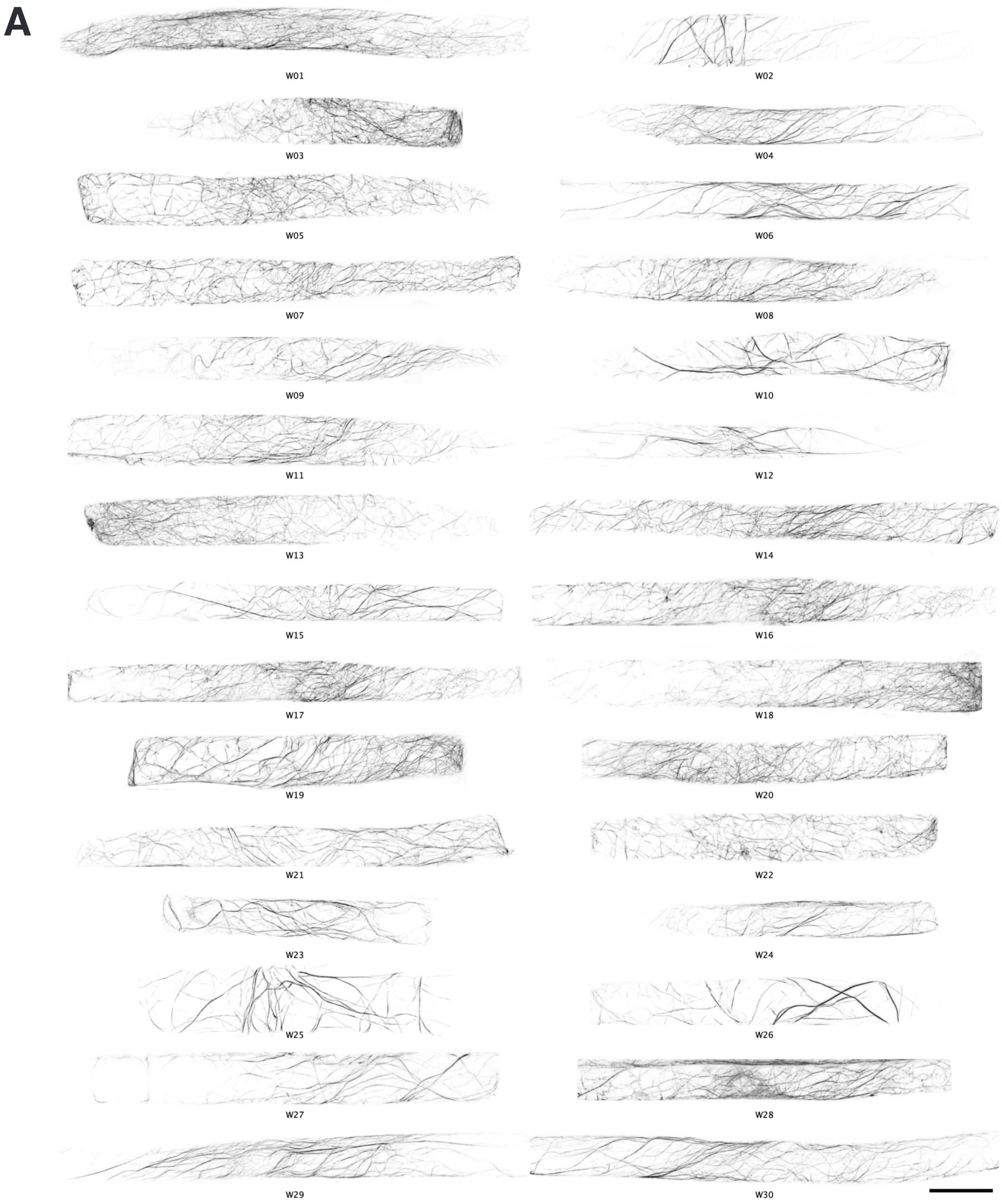
A
W01
W02
W03
W04
W05
W06
W07
W08
W09
W10
W11
W12
W13
W14
W15
W16
W17
W18
W19
W20
W21
W22
W23
W24
W25
W26
W27
W28
W29
W30

B

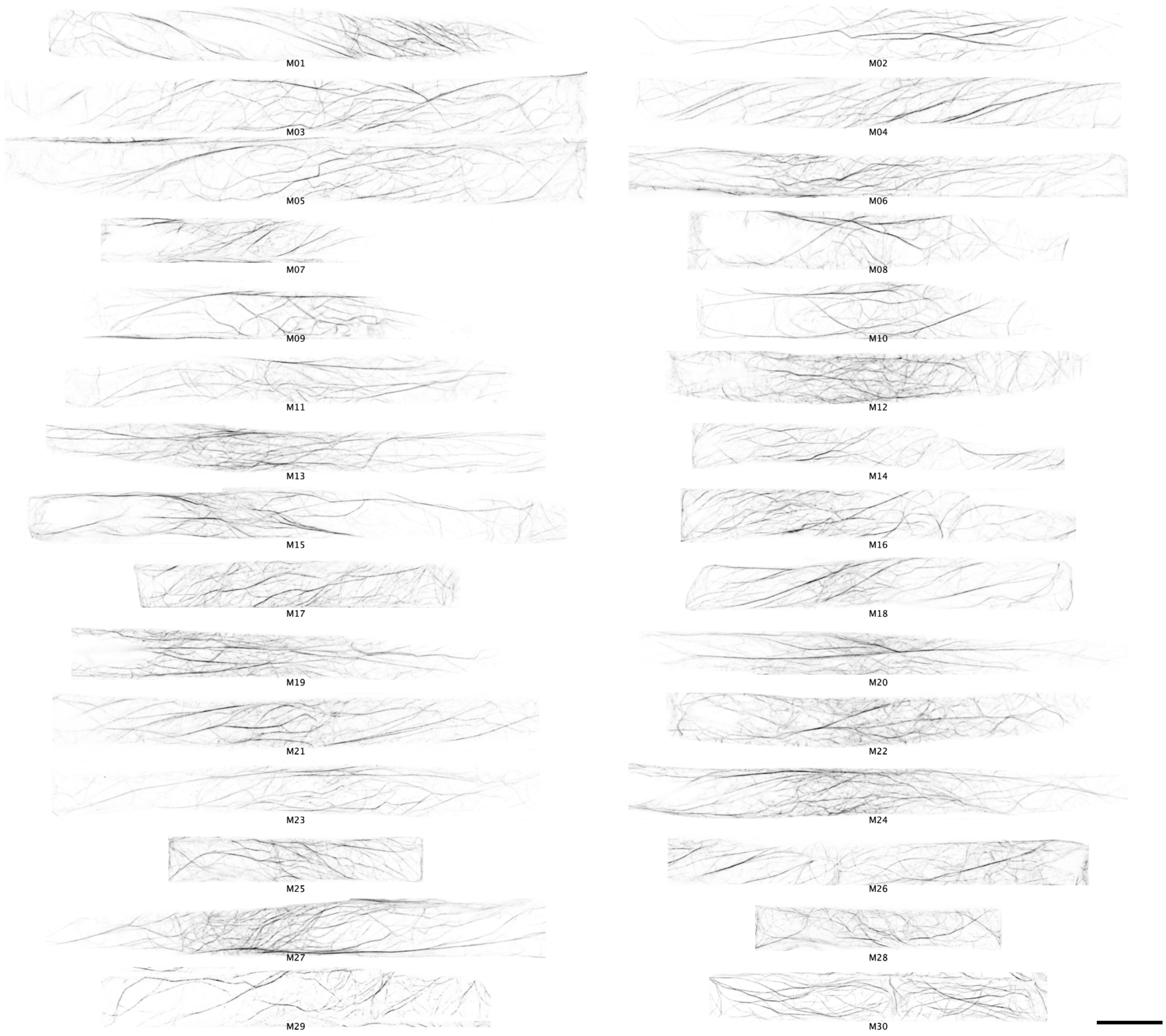


**Figure S5.** Cortical actin networks of (A) wild-type and (B) *myo11e* mutant root atrichoblast epidermal cells. Images represent maximum intensity projections of confocal stacks capturing the outer 4 to 7 μm of each cell. Fluorescent images have been inverted so that filaments appear dark on a bright background. Scale bars represent 10 μm.

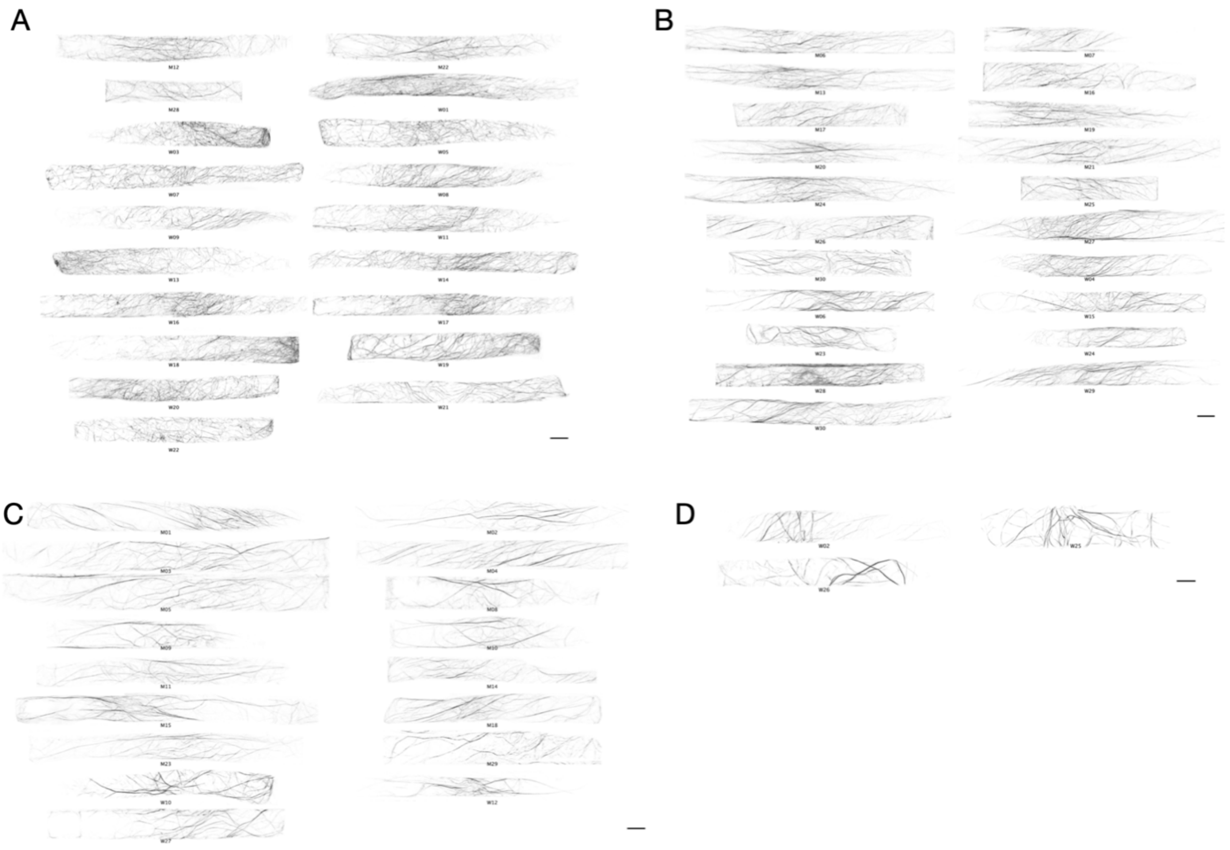


**Figure S6.** K-means clustering of cells (k = 4). (A) Cluster 1 contained cells with fine meshwork and was mostly associated with wild-type cells. (B) Cluster 2 had cells with a mixed network phenotype and contained both a fine meshwork and more prominent bundles. Both wild-type and mutant cells were found in this cluster. (C) Cluster 3 contained cells with prominent bundles and was mostly associated with mutant cells. (D) Cluster 4 contained a few cells with very prominent bundles, few fine filaments, and large filament-free areas. Scale bars are 10 µm.

**Table S1**. Pairwise correlations between different morphometric parameters for simulated networks. Positive correlations are indicated by shades of red. Negative correlations are indicated by shades of blue.

| | Occupancy | Distance | Angular Variation | Order Parameter | Bundle Parameter | Mean Weighted Angle |
|---|---|---|---|---|---|---|
| Occupancy | 1.000 | -0.931 | 0.972 | -0.960 | -0.987 | -0.047 |
| Distance | -0.931 | 1.000 | -0.882 | 0.848 | 0.911 | 0.026 |
| Angular Variation | 0.972 | -0.882 | 1.000 | -0.996 | -0.948 | -0.050 |
| Order Parameter | -0.960 | 0.848 | -0.996 | 1.000 | 0.938 | 0.050 |
| Bundle Parameter | -0.987 | 0.911 | -0.948 | 0.938 | 1.000 | 0.055 |
| Mean Weighted Angle | -0.047 | 0.026 | -0.050 | 0.050 | 0.055 | 1.000 |

**Table S2**. Pairwise correlations between different morphometric parameters for root epidermis cells. Positive correlations are indicated by shades of red. Negative correlations are indicated by shades of blue.

| | Occupancy | Distance | Angular Variation | Order Parameter | Bundle Parameter | Mean Weighted Angle |
|---|---|---|---|---|---|---|
| Occupancy | 1.000 | -0.895 | 0.196 | -0.252 | -0.900 | 0.005 |
| Distance | -0.895 | 1.000 | -0.018 | 0.095 | 0.964 | -0.042 |
| Angular Variation | 0.196 | -0.018 | 1.000 | -0.969 | -0.104 | -0.029 |
| Order Parameter | -0.252 | 0.095 | -0.969 | 1.000 | 0.189 | -0.074 |
| Bundle Parameter | -0.900 | 0.964 | -0.104 | 0.189 | 1.000 | -0.036 |
| Mean Weighted Angle | 0.005 | -0.042 | -0.029 | -0.074 | -0.036 | 1.000 |
